# Mechanism of commitment to a mating partner in *Saccharomyces cerevisiae*

**DOI:** 10.1101/2022.02.10.479952

**Authors:** Katherine C. Jacobs, Olivia Gorman, Daniel J. Lew

**Affiliations:** Department of Pharmacology and Cancer Biology, Duke University, Durham, NC 27708

## Abstract

Many cells detect and follow gradients of chemical signals to perform their functions. Yeast cells use gradients of extracellular pheromones to locate mating partners, providing a tractable model to understand how cells decode the spatial information in gradients. To mate, yeast cells must orient polarity toward the mating partner. Polarity sites are mobile, exploring the cell cortex until they reach the proper position, where they stop moving and “commit” to the partner. A simple model to explain commitment posits that a high concentration of pheromone is only detected upon alignment of partner cells’ polarity sites, and causes polarity site movement to stop. Here we explore how yeast cells respond to partners that make different amounts of pheromone. Commitment was surprisingly robust to varying pheromone levels, ruling out the simple model. We also tested whether adaptive pathways were responsible for the robustness of commitment, but our results show that cells lacking those pathways were still able to accommodate changes in pheromone. To explain this robustness, we suggest that the steep pheromone gradients near each mating partner’s polarity site trap the polarity site in place. This mechanism has evolutionary implications because it enables sexual selection for cells making more pheromone.

## Introduction

Many cells decode the spatial distribution of informative chemicals in their environment to navigate towards a destination. For example, the social amoeba *Dictyostelium discoideum* migrates up a gradient of cAMP to aggregate and form a fruiting body, neutrophils migrate up gradients of bacterial peptides or chemokines to reach sites of infection or injury, and axonal growth cones follow guidance cues to reach synaptic targets in the nervous system (Xiang et al., 2002; von Philipsborn and Bastmeyer, 2007; Devreotes et al., 2017). In many cases, the chemical signals are detected by G protein coupled receptors (GPCRs) on the cell surface, which promote reorganization of the cytoskeleton to direct movement or growth up the gradient (McCudden et al., 2005).

Cells can integrate information from several chemical and physical cues, detected by distinct sensing pathways. Due to this complexity, as well as extensive redundancy in the signaling networks that link receptor signaling to the cytoskeleton, we lack a detailed molecular understanding of how spatial gradients of signaling molecules lead to directed growth or movement (Insall, 2013). The genetically tractable budding yeast *Saccharomyces cerevisiae* provides several advantages in this regard, because it has a simplified signaling pathway with much less redundancy and appears to care about only one chemical cue: the pheromones secreted by potential mates (Ghose et al., 2022).

Haploid yeast cells of the different mating types, MAT**a** or MAT*α*, signal to each other by secreting peptide pheromones that are detected by GPCRs on the surface of cells of the opposite mating type. Pheromone-receptor binding results in activation of a MAPK pathway that induces expression of mating-specific genes and causes G1 cell-cycle arrest in preparation for mating (Cross et al., 1988; Dohlman and Thorner, 2001). MAPK activation also leads to polarization of the conserved Rho-family GTPase Cdc42. Concentration of Cdc42 at a “polarity site” at the cortex leads to orientation of actin cables that direct vesicle secretion and hence growth to the polarity site (Chiou et al., 2017). Polarized secretion enables the partners to grow toward and contact each other, and then degrade the intervening cell walls at the contact site in preparation for membrane fusion. Cell wall degradation is mediated by secretion of cell wall hydrolases, but degradation anywhere except the contact site would lead to catastrophic cell lysis. Thus, proper positioning of the polarity site is essential for mating.

How is the polarity site positioned? Landmark studies identified a scaffold protein called Far1 as a central player in getting the polarity site to the right location, and suggested an appealingly simple hypothesis for how polarity site positioning might occur (Valtz et al., 1995; Butty et al., 1998; Nern and Arkowitz, 1998; Nern and Arkowitz, 1999). Far1 binds to the Cdc42-directed GEF, and also to Gβγ formed in response to pheromone receptor activation. Thus, a cell could translate a spatial gradient of pheromone (via G protein activation and Far1-GEF recruitment) into a spatial gradient of Cdc42 activation, biasing polarity establishment to occur in the direction of the pheromone source. However, although the Far1 pathway does bias polarity establishment to occur towards a mating partner, the bias is weak and the initial polarity site is frequently mis-oriented (Henderson et al., 2019; Wang et al., 2019). One possible reason is that yeast mate in crowded conditions where a cell may be surrounded by several potential mates, so that net pheromone gradients may be uninformative (McClure et al., 2018). Mating in these conditions requires a way to identify a single partner among several suitors. Recent findings in both *S. cerevisiae* and the distantly related fission yeast *Schizosaccharomyces pombe* have revealed that yeast cells employ an “exploratory polarization” strategy to achieve this goal (Fig. 1A).

**Figure 1.**
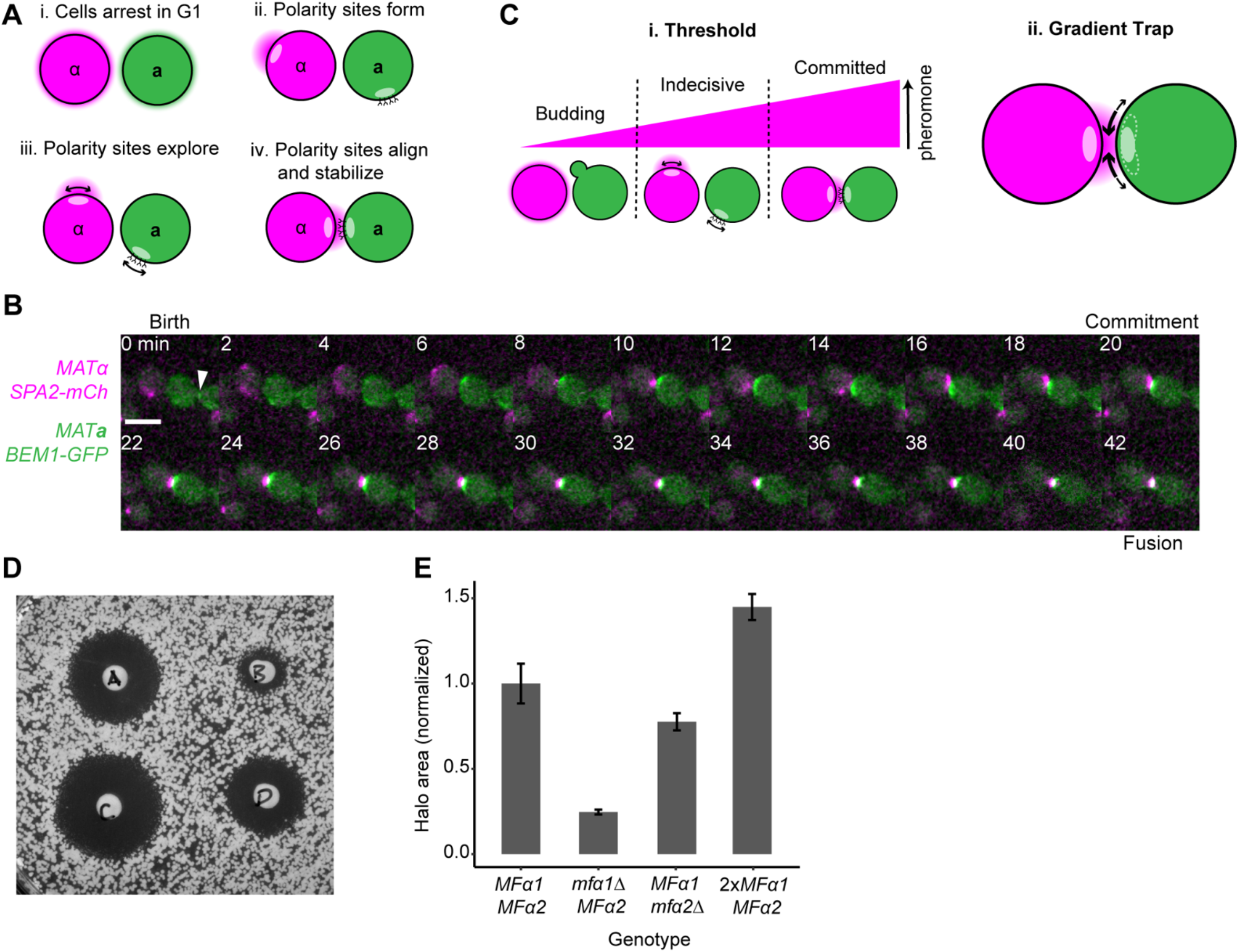
Polarity site behavior and changing α-factor secretion. (A) Exploratory polarization. Mating yeast cells secrete pheromones and respond to pheromones from the opposite mating type. Pheromone sensing leads to arrest in G1 phase (i) and polarization (ii). Pheromone secretion is focused at polarity sites, and receptors cluster at polarity sites (for simplicity, pheromone is illustrated only for α and receptors are illustrated only for **a**, but both cells secrete and sense pheromones at polarity sites). Polarity sites are then mobile (arrows, iii) until they align, at which point they stop moving (iv). (B) Localization of Bem1-GFP in a MAT**a** cell from birth (cytokinesis) to fusion. Merged 2-color maximum projection images of z-stacks from a mating mix between MAT**a** Bem1-GFP (DLY22321) and MATα Spa2-mCherry (DLY23483) with time indicated in minutes. White arrow = cytokinesis (t=0). The signal is dynamic during an “indecisive” phase until t=20 min, when it stabilizes during a “committed” phase culminating in fusion at t=42 min. Scale bar, 5 µm. (C) Threshold and gradient trap hypotheses to explain commitment. Threshold: above a low threshold concentration (dashed line) of pheromone, cells arrest in G1 and polarity sites explore the surface; above a higher threshold concentration of pheromone that is only sensed following alignment, movement stops, and polarity sites maintain position. Gradient Trap: after alignment, any further movement of the polarity site is reversed due to the steep pheromone gradient driving sites back into alignment. (D) Halo assay for α-factor secretion. Strains: A, *MFα1 MFα2* (DLY20627); B, *mfα1Δ MFα2* (DLY23250); C, 2x*MFα1 MFα2* (DLY23340); D, *MFα1 mfα2Δ* (DLY23521). (E) Quantification of halo area (normalized to *MFα1 MFα2* halo area, n = 3, error bars = standard deviation).

Following initial polarity establishment, cells relocate the polarity site in an apparent search process that can involve both lateral displacement of the polarity site along the cortex and disappearance/re-appearance of the polarity site (here we use “polarity site” to refer to a local cluster of Cdc42 and associated proteins) (Fig. 1B). Once the polarity sites in adjacent cells of opposite mating type become aligned, their position is stabilized and they stop moving (Fig. 1B). This cessation of movement constitutes commitment to a partner, and initiates the degradation of the intervening cell walls to allow fusion (Martin, 2019).

How does the polarity site “know” it should stop moving when it reaches the proper position? Previous studies proposed a mechanism whereby detection of a threshold pheromone signal is the primary trigger for commitment: we will call this the “threshold hypothesis” (Merlini et al., 2016; Clark-Cotton et al., 2021). When cells polarize, they secrete pheromone from the polarity site, signaling its location to potential partners. Pheromone receptors and G proteins become concentrated in a zone surrounding the polarity site, creating a sensitized “nose” that locally detects the pheromone secreted by potential partners (McClure et al., 2015; Merlini et al., 2016; Clark-Cotton et al., 2021). When partner polarity sites are mis-oriented, each site would detect a low pheromone concentration (Fig. 1A). However, when partners’ polarity sites are oriented toward each other, they would detect a much higher concentration of pheromone (Fig. 1A). Thus, the appealingly simple threshold hypothesis posits that there is a pheromone concentration threshold above which the polarity site is stabilized and commitment occurs (Fig. 1Ci).

Support for the threshold hypothesis comes from studies exposing yeast cells to different concentrations of synthetic pheromone. Early work found that while a low pheromone concentration was sufficient to cause cell-cycle arrest, a much higher pheromone concentration was needed to induce the growth of mating projections, which form when cells have a stably positioned polarity site (Moore, 1983). More recent work confirmed that a higher pheromone concentration was required to generate a strong polarity cluster than to arrest the cell cycle (Errede et al., 2021), and that polarity site movement was reduced as the pheromone concentration increased (Bendezú and Martin, 2013; McClure et al., 2015). However, it is not clear that cells possess the synthetic capacity to make enough pheromone to expose their partners to doses comparable to those achieved experimentally with synthetic pheromone, and other mechanisms for commitment may not require a specific pheromone threshold.

A different way to explain why polarity site movement ceases upon alignment is based on the pheromone gradients (rather than absolute pheromone concentrations) across the polarity site: we will call this the “gradient trap” hypothesis (Fig. 1Cii). This hypothesis is based on the observation that the direction of polarity site movement is biased by the Far1 pathway responding to local pheromone gradients (Ghose et al., 2021). Simulations have shown that the pheromone gradient is very steep when the polarity sites in partner cells are near each other (Clark-Cotton et al., 2021), but upon reaching alignment there would be no net gradient across the well-centered polarity site. Thus, the polarity site might be trapped in place following alignment because if it strays a little in any direction, it would be exposed to a steep gradient and its movement would be strongly biased to return to alignment.

Here we present an analysis of the robustness with which cells are able to sense polarity site alignment and cease polarity site movement when confronted with partners that secrete different amounts of pheromone. Our findings suggest that the threshold hypothesis does not suffice to explain the robust and accurate commitment capacity exhibited by mating yeast. We suggest that the steepness of local pheromone gradients can trap the polarity site at the site of cell-cell contact.

## Results

### Mating with partners that make less α-factor

We began with a test of the threshold hypothesis (Fig. 1Ci), which predicts that if the pheromone secreted by one partner is insufficient to reach the threshold concentration, then the polarity site in the other partner would not stop moving even when it becomes aligned. To test this prediction, we generated yeast strains that produce decreased levels of α-factor pheromone. Mature α-factor is cleaved from precursor polypeptides encoded by *MFα1* and *MFα2* (Michaelis and Barrowman, 2012). We deleted either *MFα1* or *MFα2*, and assayed pheromone production using the “halo” assay, which detects the growth inhibition resulting from pheromone-induced G1 arrest. Equal numbers of wild-type and mutant MATα cells were spotted onto a lawn of supersensitive MAT**a** tester cells. Pheromone secreted by the MATα cells diffuses away from the spot and arrests the growth of the underlying tester cells, and the zone of growth inhibition (halo area) provides an indication of the amount of pheromone secreted. Halo area was reduced by ∼80% for the *mfα1Δ* strain and by ∼20% for the *mfα2Δ* strain, as compared to wild-type (Fig. 1D,E). This is consistent with biochemical estimates that *MFα1* accounts for 80% of pheromone production and *MFα2* accounts for 20% (Rogers et al., 2012).

We mixed MATα cells making wild-type or decreased levels of pheromone (magenta) with wild-type MAT**a** partners (green), and imaged mating mixes. As previously reported, wild-type partners exhibited unstable and mobile polarity sites during an “indecisive” search phase followed by a “committed” phase during which partner cells’ polarity sites were stably aligned, leading to fusion (Fig. 1B). To measure how well cells commit, the behavior of each MAT**a** cell with an available MATα partner was scored as (i) successful commitment, (ii) budding (failure of G1 arrest), or (iii) continued G1 arrest with failure to commit by the end of the 120 min movie (Fig. 2A). Our expectation was that MATα cells making very little pheromone (*mfα1Δ*) would be unable to maintain G1 arrest in their partners, while partners of cells making intermediate levels of pheromone (*mfα2Δ*) might maintain G1 arrest but fail to commit. As expected, cells making less pheromone failed to efficiently arrest their partners in G1, leading to increased budding (Fig. 2A,B). Surprisingly, however, many of the cells that remained arrested did successfully commit and mate, even with *mfα1Δ* partners (Fig. 2A). This subset of cells displayed an indecisive phase of normal duration (Fig. 2C). These results are consistent with previous reports that cells secreting reduced pheromone mate less well (Brizzio et al., 1996; Huberman and Murray, 2013).

**Figure 2.**
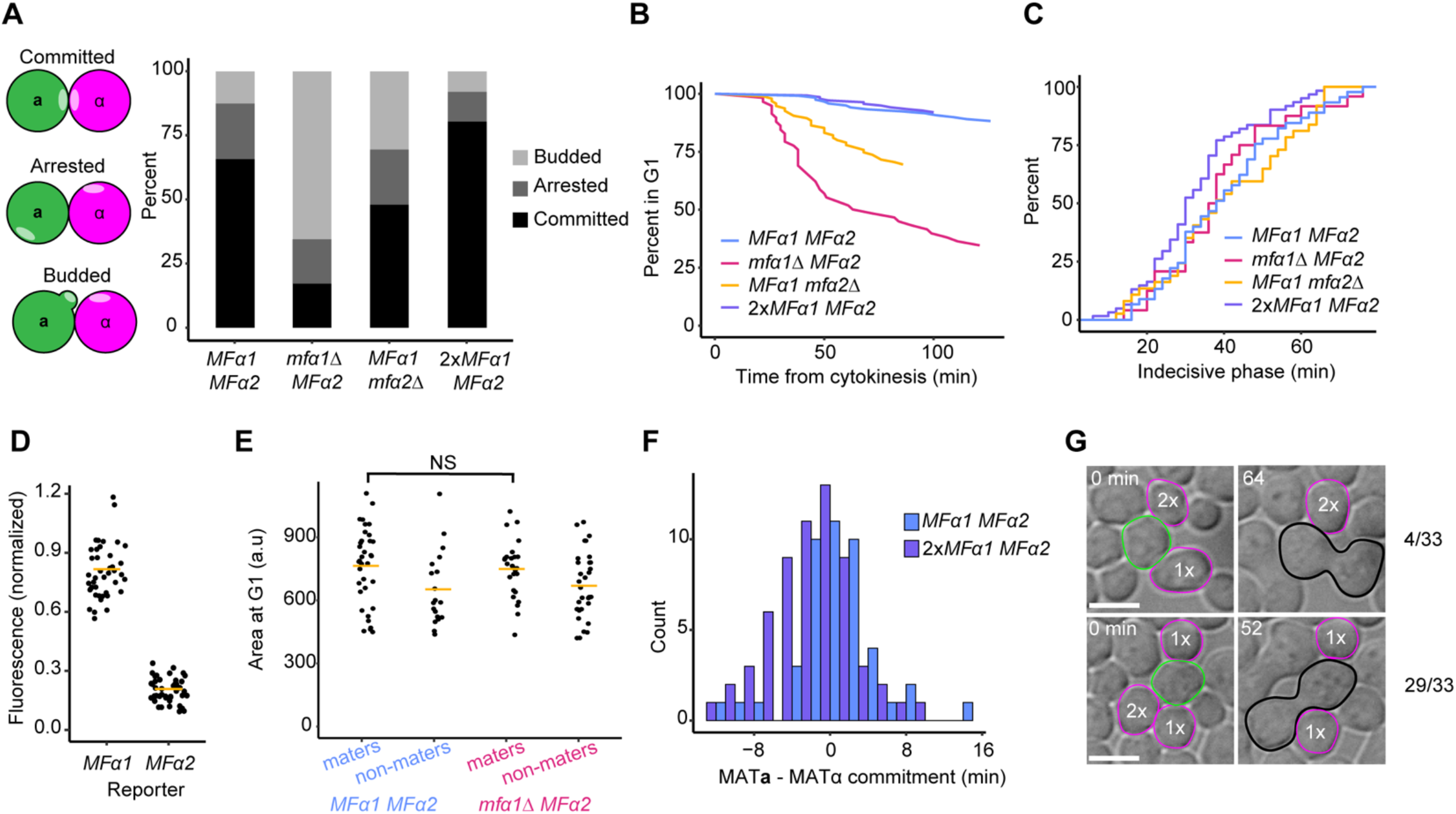
Effect of changing α-factor secretion. (A) Left: Potential outcomes for MAT**a** cells in a mating mixture after 120 minutes of co-incubation. Right: Percent of MAT**a** cells (DLY22321) with each outcome in crosses with *MFα1 MFα2* (DLY23483), *mfα1Δ MFα2* (DLY23836), *MFα1 mfα2Δ* (DLY23435), or 2x*MFα1 MFα2* (DLY23345) partners (n ≥ 58 cells). (B) Percent of cells arrested in G1 as a function of time (t_0_ = cytokinesis for each cell, G1 ends at bud emergence) in the same crosses. (C) Cumulative distribution of the duration of the indecisive phase for successful mating pairs in the same crosses (n ≥ 24 cells) (*MFα1 MFα2* v. *2xMFα1 MFα2* Kolmogorov-Smirnov (KS) test p = 0.05, all other comparisons NS). (D) Quantification of GFP fluorescence from *MFα1* (DLY23902) and *MFα2* (DLY23895) transcriptional reporters (promoter driving sfGFP) in mating mixes with DLY8502, measured at the time of commitment and normalized (*MFα1* mean set to 0.8, *MFα2* mean set to 0.2). (E) Cell area of cells in G1 phase that did or did not succeed in mating. Crosses with the indicated genotypes as in (A) (*MFα1 MFα2* maters v. *mfα1Δ MFα2* maters T test = NS). (F) Histogram of the time interval between when the MAT**a** and MATα partners commit for crosses with strains secreting wild-type or excess α-factor (n ≥ 45 pairs, crosses as in A) (KS test = NS). (G) DIC images of mating events (time, min) for a three-way mating mix where MAT**a** cells (green, DLY22321) have a choice of MATα partners (magenta) producing wild-type (1x = 1 copy of *MFα1*, DLY17505) or excess pheromone (2x = 2 copies of *MFα1*, DLY23345). Scale bar, 5 µm.

Our findings suggest that cells making too little pheromone to effectively arrest their partners are nevertheless able to stabilize the partner’s polarity site in a fraction of cells. This would seem inconsistent with the threshold hypothesis, which posits that considerably higher pheromone levels are required to stabilize polarity than to arrest the partner in G1 (Fig. 1Ci). However, a caveat is that large cell-to-cell variation in pheromone production could allow a subset of mutant cells to make wild-type levels of pheromone, enabling them to mate. Transcriptional reporters for the *MFα1* and *MFα2* promoters did not exhibit sufficient variability to sustain this interpretation (Fig. 2D). Another caveat is that the successful maters among *mfα1Δ* mutants might be the largest cells, whose increased protein synthesis capacity might allow them to make more pheromone. However, there was no significant difference in the size of maters between mutant and wild-type mating mixes (Fig. 2E). Thus, it would appear that all *mfα1Δ* mutants secrete low levels of pheromone, capable of inducing only transient cell-cycle arrest in neighboring partners. Nevertheless, those partners that do arrest manage to mate at quite high efficiency, suggesting that these low levels of pheromone suffice to halt the movement of the polarity site.

### Mating with partners that make more α-factor

The threshold hypothesis also predicts that if a cell secretes more pheromone than wild-type, then it would cause its partner to stop moving the polarity site before reaching alignment. To test this prediction, we generated a strain with an additional copy of *MFα1*, which resulted in a ∼50% increase in halo area (Fig. 1D,E). This strain exhibited even more efficient cell-cycle arrest and mating than the wild-type (Fig. 2A,B), with comparable duration of the indecisive phase (Fig. 2C). This is consistent with previous reports that strains making excess pheromone mate well (Jackson and Hartwell, 1990a; Huberman and Murray, 2013). If increased α-factor had caused premature commitment by the MAT**a** partner, we would expect to see a consistent order of commitment, with the MAT**a** partner consistently committing before the MATα partner. However, we did not find a significant difference in commitment times of the MAT**a** vs. MATα mating partners when pheromone secretion was increased (Fig. 2F). We conclude that this level of increased pheromone secretion does not cause premature commitment.

A previous study found that when MAT**a** cells are given a choice between two mating partners making different amounts of α-factor, they choose the mating partner making the highest amount (Jackson and Hartwell, 1990b). However, this study only compared MATα strains making wild-type and reduced levels of α-factor. To ask if MAT**a** cells prefer partners making wild-type or increased levels of α-factor, we imaged mating mixtures with two potential MATα partners making either wild-type or increased α-factor. We identified MAT**a** cells that had at least one potential mating partner of each genotype and scored which partner the MAT**a** cell mated with. Of 33 MAT**a** cells observed, 29 mated with a partner secreting more α-factor (Fig. 2G). Of 21 MAT**a** cells that were in contact with equal numbers of each partner, 18 mated with a partner making increased α-factor. We conclude that MAT**a** cells can discriminate between MATα cells making wild-type and increased levels of pheromone, and that they choose the partner making the most pheromone.

### Effect of the Bar1 α-factor protease

Interpretation of the experiments discussed above depends on the key assumption that a change in the level of pheromone produced/secreted by the cells of one mating type leads to a change in the level of pheromone perceived by the cells of the other mating type. However, this need not always be the case: MAT**a** cells produce and secrete a protease called Bar1 that degrades α-factor, which has been proposed to sculpt the pheromone landscape in various ways (Barkai et al., 1998; Jin et al., 2011; Rappaport and Barkai, 2012). Consistent with previous reports (Chan and Otte, 1982; Conlon et al., 2016), we found that MAT**a** *bar1Δ* cells had a very low (2%) mating efficiency in our slab mating conditions (Fig. 3B). MAT**a** *bar1Δ* cells should experience higher pheromone levels and, thus, commit prematurely with incorrect orientation. As expected, most MAT**a** *bar1Δ* cells stabilized their polarity sites at random locations and not towards potential partners (Fig. 3A). Thus, Bar1 is necessary to prevent premature (incorrect) commitment by MAT**a** cells.

**Figure 3.**
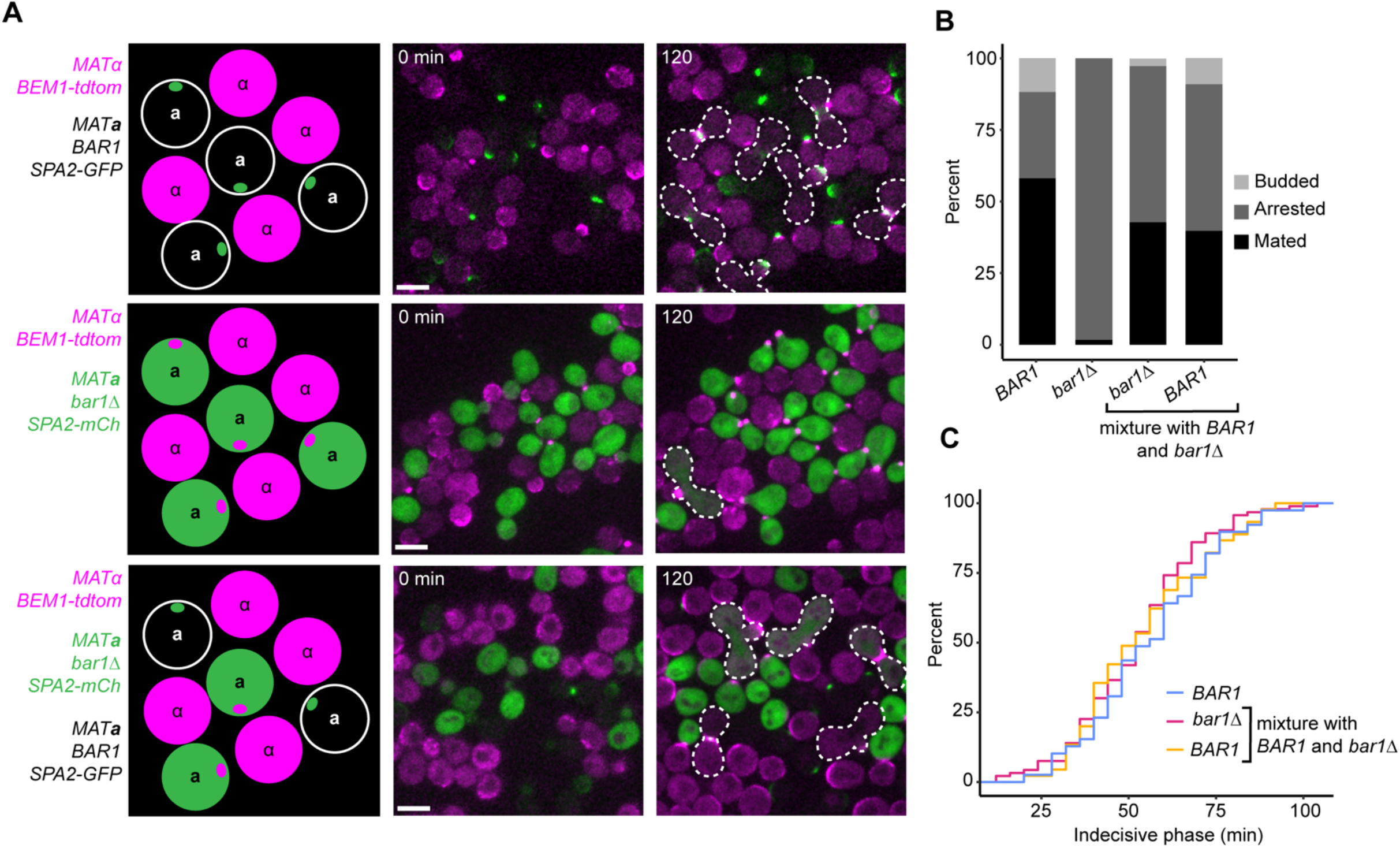
Secreted Bar1 acts globally to avoid saturation with α-factor. (A) Mating mixes between wild-type cells (top, DLY21840 x DLY12944), MAT**a** *bar1Δ* x MATα (middle, DLY23430 x DLY12944), and three-way mix with MAT**a** *BAR1* and MAT**a** *bar1Δ* x MATα (middle, DLY23430 and DLY21840 x DLY12944) with probes indicated in the left panels were imaged. Representative maximum-projection images from z-stacks of mating mixtures are shown at 0 min and 120 min. Zygotes are outlined in white dashed lines. The last panel of the middle row shows randomly oriented mating projections (shmoos) formed by MAT**a** *bar1Δ* cells. Scale bar, 5 µm. (B) Percent of MAT**a** cells that mated, budded, or remained arrested in the same mating mixtures (n ≥ 136 cells). (C) Cumulative distributions of the duration of the indecisive phase for mating events in the same mixtures (except for the MAT**a** *bar1Δ* x MATα cross which produced insufficient successful matings to score) (n ≥ 39 cells) (all KS tests = NS).

Bar1 is secreted from the polarity site and some fraction of the secreted Bar1 remains attached to the cell wall (Moukadiri et al., 1999). Thus, Bar1 could act locally near the polarity site to lower local pheromone levels, ensuring that above-threshold concentrations sufficient to induce commitment do not occur until polarity sites are aligned. Alternatively, secreted Bar1 could be dispersed rapidly and act globally to lower α-factor concentration on the slab. If Bar1 acts globally, then Bar1 secreted by wild-type “helper” cells should rescue the mating defect of *bar1Δ* cells on the same slab. But if Bar1 specifically helps the cell that secretes it, then wild-type helper cells would not rescue the mating defect of *bar1Δ* cells. In our mating conditions, wild-type helper cells fully rescued the mating defect of *bar1Δ* cells (Fig. 3A-C), consistent with a previous report (Conlon et al., 2016). This finding suggests that Bar1’s primary role is to globally lower the α-factor concentration to prevent premature commitment.

*BAR1* expression increases in response to perceived α-factor (Manney, 1983; Aymoz et al., 2018). Thus, adjustable Bar1 expression could buffer the α-factor concentration around some optimal value (Fig. 4A). If that were the case, then the finding that cells mate well when α-factor secretion rate is altered might simply reflect compensatory alteration in the secretion of Bar1. To test whether pheromone-regulated *BAR1* expression is important for cells to adjust to their environment, we replaced the endogenous *BAR1* promoter with the constitutive *TDH3* promoter. For these experiments, we altered the pheromone level by altering the ratio of MAT**a** to MATα partners. We mixed cells at ratios varying over two orders of magnitude, between 10:1 and 1:10 (MATα:MAT**a**), and imaged the behavior of the MAT**a** cells that were touching MATα partners. Wild-type cells committed with similar efficiency regardless of the ratio (Fig. 4B), consistent with the findings for partners that produce different levels of pheromone (Fig. 2). We found that cells making constitutive levels of Bar1 also committed efficiently at ratios varying between 10:1 and 1:10 (Fig. 4B). We conclude that inducible Bar1 expression is not responsible for the robustness with which cells can mate in contexts with different levels of pheromone.

**Figure 4.**
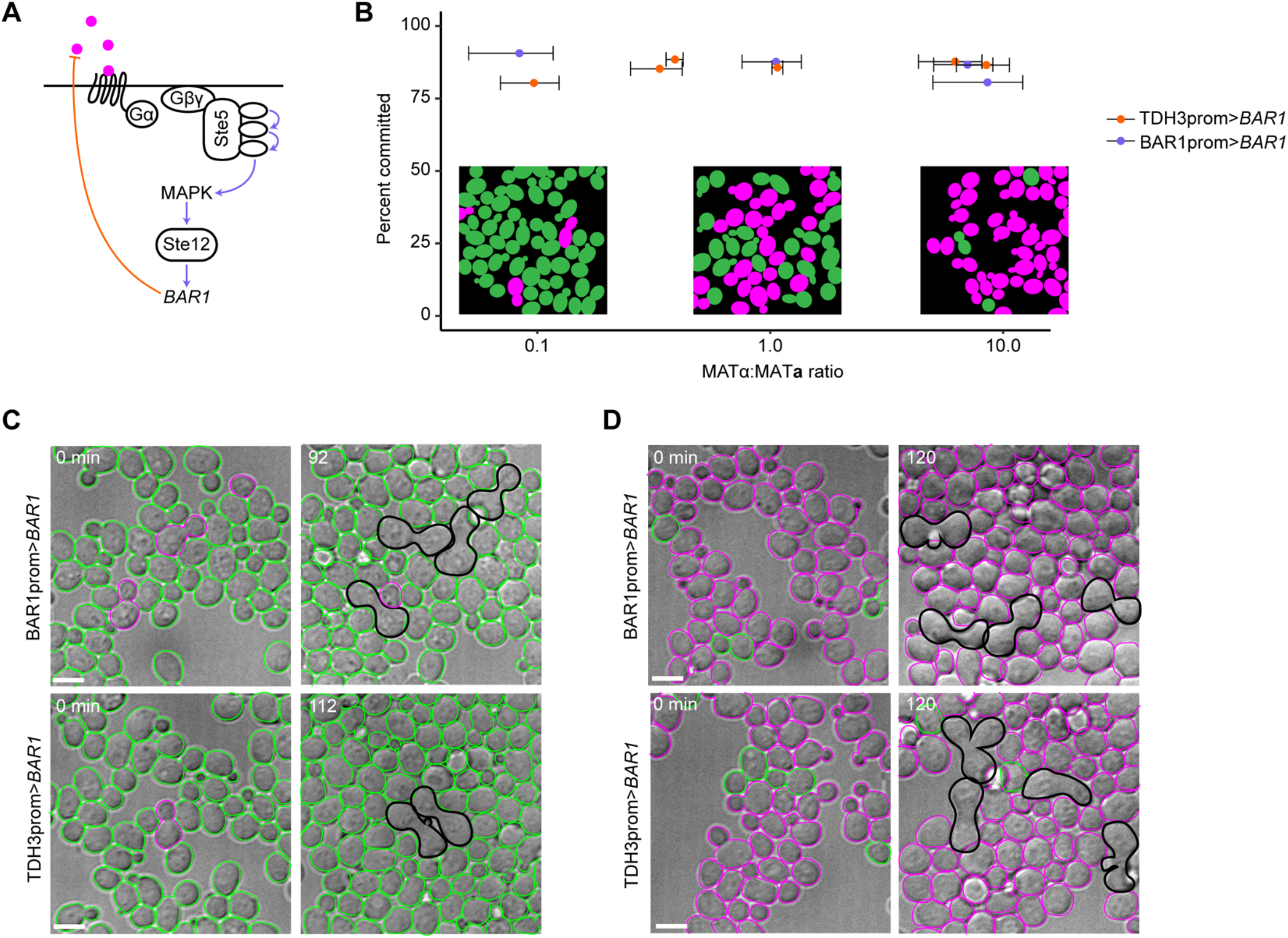
Pheromone-induced *BAR1* expression is not necessary for efficient mating in conditions with different levels of α-factor. (A) Pheromone signaling via GPCR triggers a MAPK cascade that activates the transcription factor Ste12, which stimulates expression of Bar1, potentially functioning as a homeostatic mechanism to maintain external α-factor levels. (B) Mating mixes (wild-type, DLY8502 x DLY9070, purple; constitutive-*BAR1*, DLY23827 x DLY9070, orange) were set up with different ratios of MAT**a**:MATα varying 100-fold from 1:10 to 10:1. The percent of MAT**a** cells that commit to a partner was scored in each condition (Y axis, n ≥ 50 cells). The MAT**a**:MATα ratio was calculated as the average from 8 stage positions: error bars indicate standard deviation. (C,D) Representative DIC images of mating mixtures with ∼1:10 (C) and ∼10:1 (D) MAT**a**:MATα ratios. Green, MAT**a** cells; magenta, MATα cells; black, zygotes. Scale bar, 5 µm.

### Mating without MAPK-mediated adaptive pathways

The finding that cells commit well to partners making quite different levels of pheromone appears incompatible with the hypothesis that a fixed threshold level of pheromone is required to halt polarity site movement. However, it remains possible that cells have an adjustable threshold, using an adaptive pathway to modulate the responsiveness of polarity-site movement to perceived pheromone. With an adjustable threshold, more pheromone would be required to stop polarity site movement when cells are exposed to high-pheromone environments (as with mating to pheromone overproducers) than to low-pheromone environments (as with mating to pheromone underproducers). In effect, commitment would respond to the fold-change in pheromone level that occurs during alignment rather than the absolute pheromone level. An adaptive threshold would require a negative feedback pathway that lowers polarity-site responsiveness when ambient pheromone levels rise.

There are multiple documented negative feedback pathways in the pheromone signaling circuit, all mediated by altering MAPK activity (Fig. 5A). In addition to inducing *BAR1* expression, MAPK activation induces expression of *SST2*, a regulator of G protein signaling (RGS) protein that inactivates Gα (Dohlman et al., 1996; Apanovitch et al., 1998), and *MSG5*, a phosphatase that dephosphorylates the MAPK (Doi et al., 1994; Chen and Thorner, 2007). MAPK also directly binds (Bhattacharyya et al., 2006) and phosphorylates (Repetto et al., 2018) the scaffold Ste5, both subunits of Gβγ (Li et al., 1998; Choudhury et al., 2018; Abdul-Ganiyu et al., 2021), and the upstream MAPKK (Errede and Ge, 1996), all of which appear to be adaptive pathways. Other adaptive pathways may also be triggered by MAPK activation (Yu et al., 2008).

**Figure 5.**
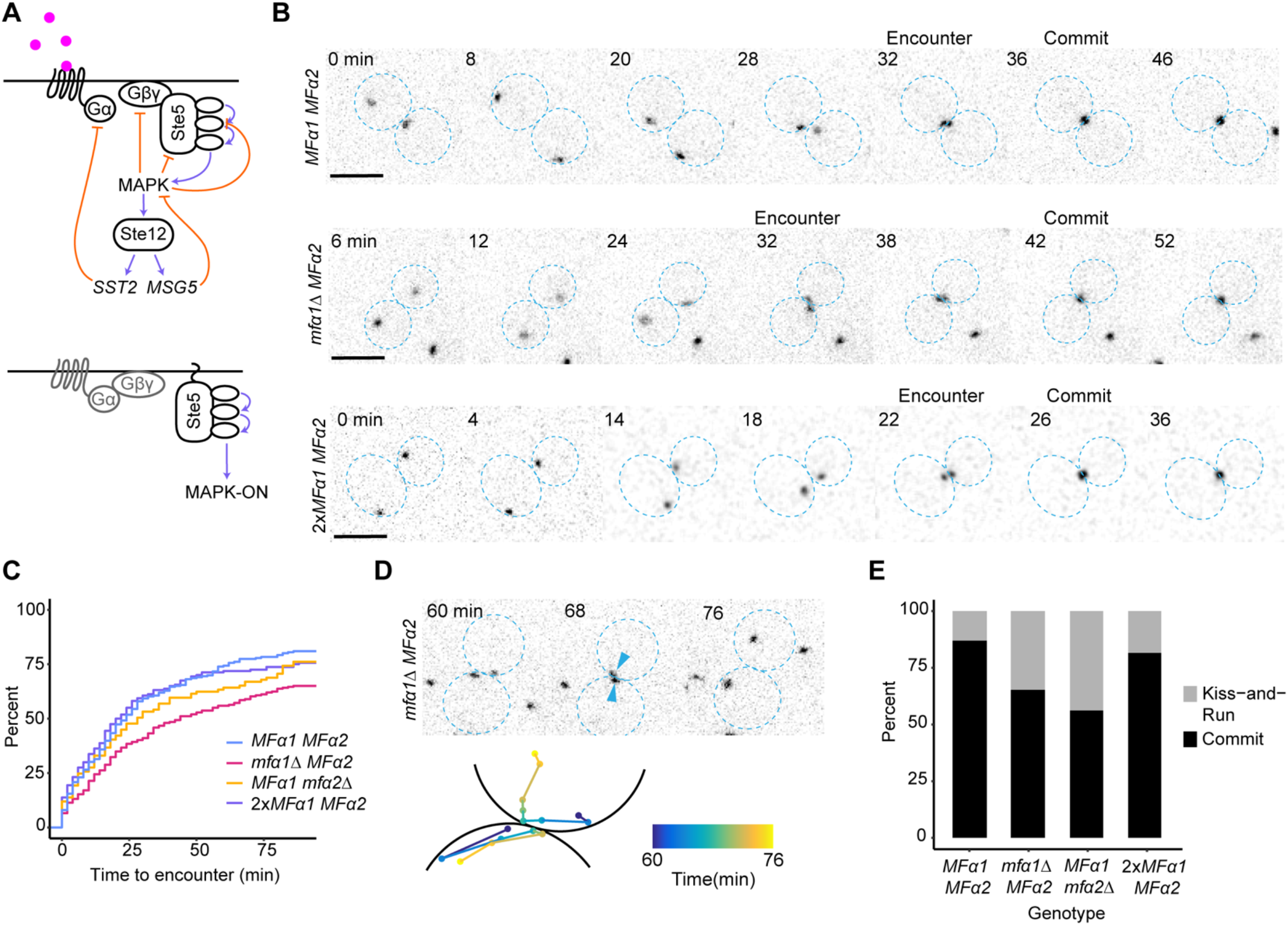
Effect of changing α-factor secretion in cells with clamped MAPK activity. (A) Top: Schematic of known negative feedback loops (orange) in the pheromone response pathway. Bottom: Ste5-CTM clamps MAPK activity at a high level that is not further induced by pheromone, thus eliminating all negative feedback via MAPK. (B) Representative inverted maximum projection images of z-stacks from mating mixes with polarity sites labeled with Spa2-mCherry in cells with clamped MAPK activity. MAT**a** cells (DLY20626) were mixed with MATα partners making wild-type levels of α-factor (top, DLY20627), low levels of α-factor (middle, DLY23250), or excess α-factor (bottom, DLY23340). Partners are outlined in blue, and the times of polarity site encounters and commitment are indicated. Scale bar, 5 µm. (C) Cumulative distributions of the time to polarity site encounter from the same movies (n ≥ 109 pairs) (*MFα1 MFα2* x *mfα1Δ MFα2* KS test p = 0.0003, all other comparisons NS). (D) Example of a kiss-and-run encounter from the mating mix with cells making low levels of α-factor. Bottom shows track of polarity site centroid for the same cells, with color indicating time: polarity sites are initially distant (dark blue), approach and almost meet (light blue) but then move further apart (yellow). (E) Percent of encounters that result in commitment or kiss-and-run (n ≥ 105 encounters) from the indicated mating mixes.

To eliminate all MAPK-mediated feedback pathways, we clamped MAPK activity at a high level by expressing a version of the MAPK scaffold protein Ste5 with a transmembrane domain (Ste5-CTM) (Fig. 5A). This results in MAPK activation even in the absence of pheromone, and there is no further MAPK activation upon exposure to pheromone (Pryciak and Huntress, 1998; McClure et al., 2015). In these strains, therefore, all adaptive pathways via the MAPK would no longer be affected by the level of pheromone in the environment, eliminating any adaptive threshold that involves MAPK.

We generated strains expressing reduced (*mfα1Δ* or *mfα2Δ*) or increased (2x *MFα1*) pheromone levels in a background where the sole copy of Ste5 was Ste5-CTM. Induction of Ste5-CTM expression caused cells to arrest in G1 and make a strong but mobile polarity site (Fig. 5B) (McClure et al., 2015; Ghose et al., 2021). We quantified how long it took polarity sites of partner cells to encounter each other in mating mixes, and asked if encounters resulted in commitment (cessation of movement). We found that polarity site encounters occur to a similar degree and on a similar timescale regardless of the amount of pheromone made by the mating partner (Fig. 5C). This suggests that the efficacy with which pheromone gradients bias the direction of polarity site movement is robust to the absolute level of pheromone, consistent with studies exposing cells to calibrated gradients of synthetic pheromone (Paliwal et al., 2007; Moore et al., 2008; Dyer et al., 2013).

Each encounter has two possible outcomes: polarity sites can become stabilized (“commit”), or polarity sites can keep moving (“kiss-and-run”) (Fig. 5D). We found that *mfα1Δ* and *mfα2Δ* mating mixtures displayed a modestly increased frequency of kiss-and-run encounters, consistent with the idea that commitment becomes less effective at low pheromone doses (Fig. 5E). However, the majority of encounters still resulted in commitment, regardless of the amount of pheromone made by the mating partner (Fig. 5E). Thus, cells that lack the ability to adjust MAPK signaling in response to pheromone can still locate and commit to mating partners when pheromone levels are altered. This is inconsistent not only with the fixed-threshold model but also with a model in which MAPK-mediated adaptive pathways produce an adjustable threshold.

### Mating with partners that make less a-factor

Our experiments thus far focused on the effects of changing α-factor secretion on the MAT**a** partner. To extend our results to **a**-factor we took a similar approach. **a**-factor is a prenylated peptide made by extensive processing from the primary products of *MFA1* and *MFA2* (Michaelis and Barrowman, 2012). However, deleting either *MFA1* or *MFA2* did not yield an appreciable change in secreted **a**-factor as measured by the halo assay (Fig. 6A), suggesting that either the halo assay is less sensitive to variation in **a**-factor secretion or that compensatory mechanisms adjust **a**-factor secretion when one gene is eliminated. We therefore deleted *STE24*, which encodes one of the enzymes involved in processing mature **a**-factor from the precursor. Consistent with a previous report (Huyer et al., 2006), the *ste24Δ* strain secreted somewhat less **a**-factor as measured by the halo assay (Fig. 6B).

**Figure 6.**
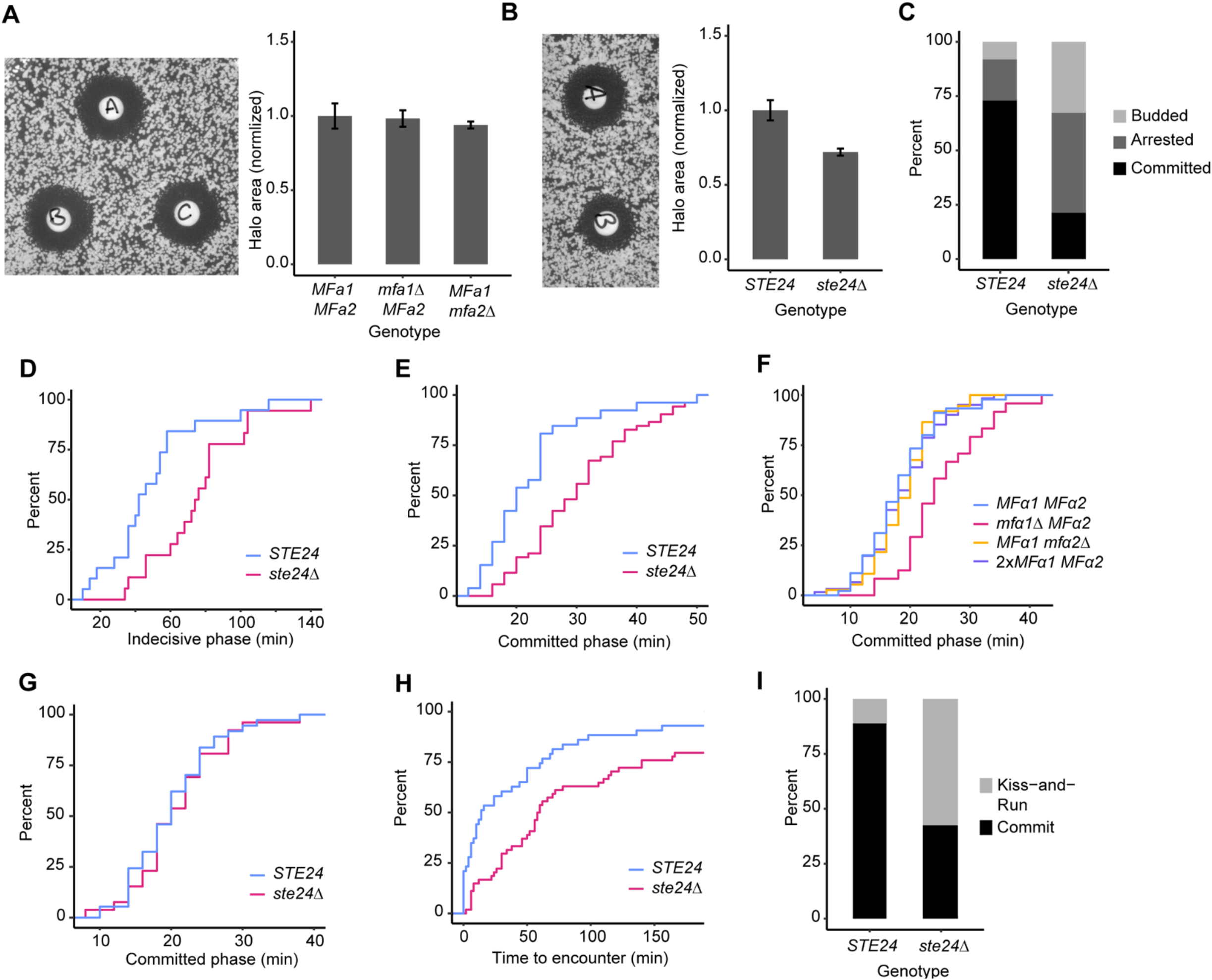
Effect of changing a-factor secretion. (A) Halo assay for **a**-factor production. Strains: A, *MFa1 MFa2* (DLY20626); B, *mfa1Δ MFa2* (DLY23990); C, *MFa1 mfa2Δ* (DLY23991). Right: Quantification of halo area (normalized to *MFa1 MFa2* halo area, n = 3, error bars = standard deviation). (B) Halo assay and quantification as in (A). Strains: A, *STE24* (DLY20626); B, *ste24Δ* (DLY24026). Right: Quantification of halo area. (C) Percent of MATα cells (DLY23483) with the indicated outcome in crosses with *STE24* (DLY22321) or *ste24Δ* (DLY24027) partners (n ≥ 37 cells). (D,E) Cumulative distributions of the duration of the indecisive (D) and committed (E) phases of successful mating events from the same mixes (n ≥ 18 cells) (indecisive phase KS test p = 0.0008, committed phase KS test p = 0.0007). (F) Cumulative distribution of the duration of the committed phase of mating events with altered α-factor (n ≥ 24 cells). Mating mixes were as in Fig. 2A. (*MFα1 MFα2* v. *mfα1Δ MFα2* KS test p = 0.001, all other comparisons, NS). (G) Cumulative distribution of the duration of the committed phase of mating events with clamped MAPK activity (n ≥ 26 cells) (KS test = NS). Strains: MAT**a** *STE24* (DLY22939), MAT**a** *ste24Δ* (DLY24026), MATα (DLY20627). (H) Cumulative distribution of the time to encounter in the same mixes as in (G) (n ≥ 43 pairs) (KS test p = 0.001). (I) Percent of encounters that result in commitment or kiss-and-run (n ≥ 50 encounters) in the same mixes as in (G).

As with cells secreting less α-factor, *ste24Δ* mutants secreting less **a**-factor induced a less potent G1 arrest in wild-type partners (Fig. 6C). Of the partners that remained arrested in G1, some were able to commit and mate, though many had not done so by the end of the movie (Fig. 6C). Even among those cells that successfully mated, the mating partners took longer to find and commit to cells secreting reduced levels of **a**-factor (Fig. 6D). These results suggest that the level of **a**-factor secretion is an important determinant of the efficiency of the search for a partner. However, once a partner is located, reduced **a**-factor secretion did not prevent commitment, suggesting that as for α-factor, a threshold concentration of **a**-factor is not responsible for stabilizing the polarity site of the MATα partner.

A previous study observed an accumulation of unfused pre-zygotes in matings where one partner secreted less pheromone (Brizzio et al., 1996). Cell fusion defects are suggestive of a prolonged committed phase, when partners are stably oriented toward each other but fail to fuse. In *ste24Δ* mutants secreting less **a**-factor, mating pairs indeed exhibited a prolonged committed phase compared to wild-type (Fig. 6E). We also observed a prolonged committed phase among mating pairs where the *mfα1Δ* mutant secreted less α-factor (Fig. 6F). Thus, the level of both pheromones affects the rate at which co-oriented committed cells can remove the intervening cell walls to achieve fusion.

Why would fusion take longer in situations with reduced pheromone? One dose-dependent output of the pheromone concentration is the degree of MAPK activation, which rises before fusion (Conlon et al., 2016) and may control the rate of cell wall degradation. To test whether the effects of pheromone level on fusion kinetics were due to modulation of MAPK activity, we generated a *ste24Δ* strain in which MAPK activity was locked at a high level using Ste5-CTM. In this context, committed mating pairs with *ste24Δ* mutants took no longer than wild-type cells to fuse (Fig. 6G). We infer that a late surge in MAPK activation is required for timely cell wall degradation.

We also measured the efficacy with which MAPK-clamped cells aligned their polarity sites when faced with partners secreting less **a**-factor. Polarity site encounters were only slightly reduced (Fig. 6H) compared to wild-type cells in this context, similar to what we had observed for encounters with cells secreting less α-factor (Fig. 5C). Although commitment still occurred, we observed a significant increase in kiss-and-run encounters (Fig. 6I). In summary, these findings with mutants secreting less **a**-factor are broadly consistent with our findings with mutants secreting less α-factor. In the main, these results do not support the predictions of fixed-threshold models for commitment. Instead, they suggest that mechanisms for locating and committing to partners are remarkably robust with regard to pheromone secretion rates.

## Discussion

### Mechanism of commitment

Previous studies had shown that exposure to high levels of synthetic pheromone could cause polarity site movement to stop (Moore et al., 2008; Dyer et al., 2013; Hegemann et al., 2015; McClure et al., 2015; Merlini et al., 2016). In combination with the recognition that both pheromone secretion and pheromone sensing occurs at polarity sites (McClure et al., 2015; Merlini et al., 2016; Clark-Cotton et al., 2021), a simple and plausible hypothesis was that sensing a pheromone concentration above some pre-set threshold causes movement to stop (the threshold hypothesis). However, it was not clear that cells possessed the pheromone synthesis capacity to expose their partners to the pheromone doses employed in experiments with synthetic pheromone. Cells are thought to produce up to 1400 molecules of α-factor per second (Rogers et al., 2012), and even if all of those were secreted from one partner’s polarity site, the polarity site in an adjacent partner would only sense about 4 nM pheromone (Clark-Cotton et al., 2021), well below the doses of synthetic pheromone required to halt movement. Moreover, using a pre-set threshold is a fragile mechanism in that there would be a narrow optimum level of pheromone secretion: too little and the partner would never sense enough pheromone to commit; too much and the partner would commit prematurely in the wrong location. We found that cells committed accurately to partners that secreted only ∼20% as much pheromone as wild-type (which was insufficient to produce a prolonged cell-cycle arrest) as well as to partners secreting excess pheromone. These and other findings argue against the simple threshold hypothesis.

We considered two other mechanisms that could explain why polarity sites in partner cells commit (stop moving) when they reach alignment: (i) sensing some fold-increase in pheromone concentration causes movement to stop (the adaptive threshold hypothesis); and (ii) steep pheromone gradients trap the moving polarity site upon reaching alignment (the gradient trap hypothesis). We note that these ideas are not mutually exclusive.

The adaptive threshold hypothesis and the gradient trap hypotheses can both explain why pheromone sensing is needed for commitment but there is no narrow optimum level of pheromone secretion. No matter what level of pheromone is secreted by a cell, it will be the case that pheromone levels detected at a polarity site increase significantly upon alignment, and that pheromone gradients are steep for polarity sites slightly out of alignment. These mechanisms could therefore produce the robust commitment that we documented.

The pheromone signaling pathway is very well studied and many potential adaptive pathways have been identified, but all known adaptive pathways rely on modulating MAPK activity in response to pheromone levels. By generating strains with MAPK activity clamped at a high level, we were able to show that modulation of MAPK in response to pheromone is not required for robust commitment. Thus, with the caveat that unknown adaptive pathways not operating through the MAPK may yet be discovered, we suggest that the gradient trap hypothesis is most likely to explain the robustness of commitment.

### Advantages of gradient-mediated commitment

We found that gradient-mediated guidance of polarity site movement led to polarity-site encounters with similar efficiency over a wide range of pheromone secretion rates. The robustness of such biased polarity site movement is consistent with studies of yeast cells exposed to artificial pheromone gradients, which found that yeast cells can orient polarity up-gradient over a wide range of mean pheromone levels, extending to levels well above the receptor K_D_ (Segall, 1993; Paliwal et al., 2007; Moore et al., 2008; Dyer et al., 2013). Exploiting gradients to promote commitment means that commitment will only occur upon alignment of the partner polarity sites.

The rate of pheromone production varies among yeast strains derived from wild isolates (McClure et al., 2018), and pheromone production incurs a fitness cost (Smith and Greig, 2010). If the level of pheromone production is an indicator of the fitness of a potential mate, that may enable sexual selection driving yeast to produce more pheromone than would be needed in order to mate (analogous to the fitness advertising value of a peacock’s tail) (Huberman and Murray, 2013). Consistent with this idea, we found that MAT**a** cells given a choice of potential partners making normal or excess levels of pheromone disproportionately chose the partner making the most pheromone. Thus, pheromone gradient-based guidance and commitment mechanisms would allow cells to select the fittest potential mates when choosing among multiple potential mating partners.

### Elevated MAPK promotes rapid cell wall removal at the contact site following commitment

Although commitment occurred with partners producing low levels of pheromone (*mfα1Δ* and *ste24Δ* cells), such pairings took longer to fuse (measuring from the time of initial commitment). This finding is consistent with a previous report (Brizzio et al., 1996), and suggests that high levels of both pheromones are needed to promote efficient degradation of the intervening cell walls. MAPK activity rises shortly before fusion (Conlon et al., 2016), and the absolute pheromone level may determine the degree of MAPK activation in committed cells. We found that inducing high MAPK activity independent of pheromone rescued the cell fusion defect of *ste24Δ* mating pairs, suggesting that the MAPK activation level controls the rate of cell wall degradation at the contact site and hence the time that cells need to fuse following commitment. Concentrated pheromone release leading to high MAPK activity is also thought to trigger fusion in *S. pombe* (Dudin et al., 2016).

### Role of extracellular α-factor degradation

Like secretion of pheromone, secretion of the α-factor protease Bar1 would presumably occur from the polarity site. This raised the possibility that under conditions that permit the development of pheromone gradients, Bar1 secreted by each cell might primarily affect the pheromone gradients detected by that cell. In theory, that could enhance discrimination among potential partners in crowded environments (Jin et al., 2011; Rappaport and Barkai, 2012). Alternatively, secreted Bar1 might constitute a shared resource for the population of mating cells, acting to prevent a global increase in pheromone levels. A global role for secreted Bar1 was proposed for cells mating in liquid environments (Banderas et al., 2016), where liquid flows would tend to yield well-mixed pheromone and Bar1. The mean pheromone concentration would then reflect the ratio between MATα (pheromone-producing) and MAT**a** (Bar1-producing) cells, allowing cells to assess whether it was worth their while to arrest and look for mates or not, taking into account both the availability of partners secreting pheromone and competitors secreting Bar1.

We found that in our experimental conditions (i.e. on solid slabs with limited volume and high cell densities), Bar1 secreted by wild-type cells rescued the mating defect of neighboring *bar1Δ* cells, which suggests that secreted Bar1 acts globally to lower the α-factor concentration to prevent premature commitment. Bar1 synthesis is induced in response to pheromone (Manney, 1983; Aymoz et al., 2018), raising the possibility that Bar1 could act homeostatically to keep external pheromone levels around some optimum setpoint. However, we found using promoter-swapped strains that cells constitutively making the same amount of Bar1 regardless of pheromone were just as good at mating, over a wide range of partner cell densities, as wild-type cells. We conclude that under our mating conditions, the primary role of Bar1 is simply to prevent a build-up of pheromone in the environment.

### Effects of reducing secretion of a-factor versus α-factor

The difference in chemical properties of prenylated **a**-factor versus unprenylated α-factor peptides has led to the suggestion that they might differ in the information they convey to the partner (Michaelis and Barrowman, 2012; Anders et al., 2021). We found that reducing the level of pheromone secretion had similar qualitative effects for either pheromone, with mutant strains inducing (i) less prolonged cell cycle arrest in the partner, (ii) a less efficient search process to align polarity sites (delays in polarity site encounters), (iii) less effective commitment (more frequent kiss-and-run encounters), and (iv) delayed fusion. However, there were quantitative differences: *ste24Δ* mutants secreting less **a**-factor led to more severe defects in partner search and commitment than *mfα1Δ* mutants secreting less α-factor, whereas *mfα1Δ* mutants led to more severe defects in G1 arrest of the partner than *ste24Δ* mutants. Because of differences in the halo assays used to assess the rates of pheromone secretion (see Methods), we cannot directly compare rates of **a**-factor versus α-factor secretion in our mutants. Nevertheless, these differences suggest that it may take different levels of **a**-factor versus α-factor to induce specific behaviors in their partners. It will be interesting to determine whether each pheromone is used in similar or different ways to convey information to the mating partner.

## Methods

### Yeast strains

Yeast strains were constructed with standard molecular genetic techniques, and are listed in Table 1. All strains were generated in the YEF473 background (*his3-Δ200 leu2-Δ1 lys2-801 trp1-Δ63 ura3-52*), with the exception of DLY15660 which is in the 15D background (*ade1 his2 leu2-3,112 trp1-1 ura3Δns*). The following alleles were previously described: *BEM1-GFP:LEU2* (Kozubowski et al., 2008), *BEM1-tdTomato:HIS3* (Howell et al., 2012), *SPA2-mCherry:kan*^*R*^ (Howell et al., 2009; Woods et al., 2015), *SPA2-mCherry:hyg*^*R*^ (McClure et al., 2015), *rsr1::HIS3* (Schenkman et al., 2002), *WHI5-tdTomato:URA3* (Moran et al., 2019), *WHI5-GFP:HIS5* and *sst2::URA3* (Clark-Cotton et al., 2021), *bar1::URA3, SPA2-GFP:HIS3* and *ste5:P*_*GAL1*_*-STE5-CTM:P*_*ADH1*_*-GAL4BD-hER-VP16:LEU2* (Henderson et al., 2019).

**Table 1.**
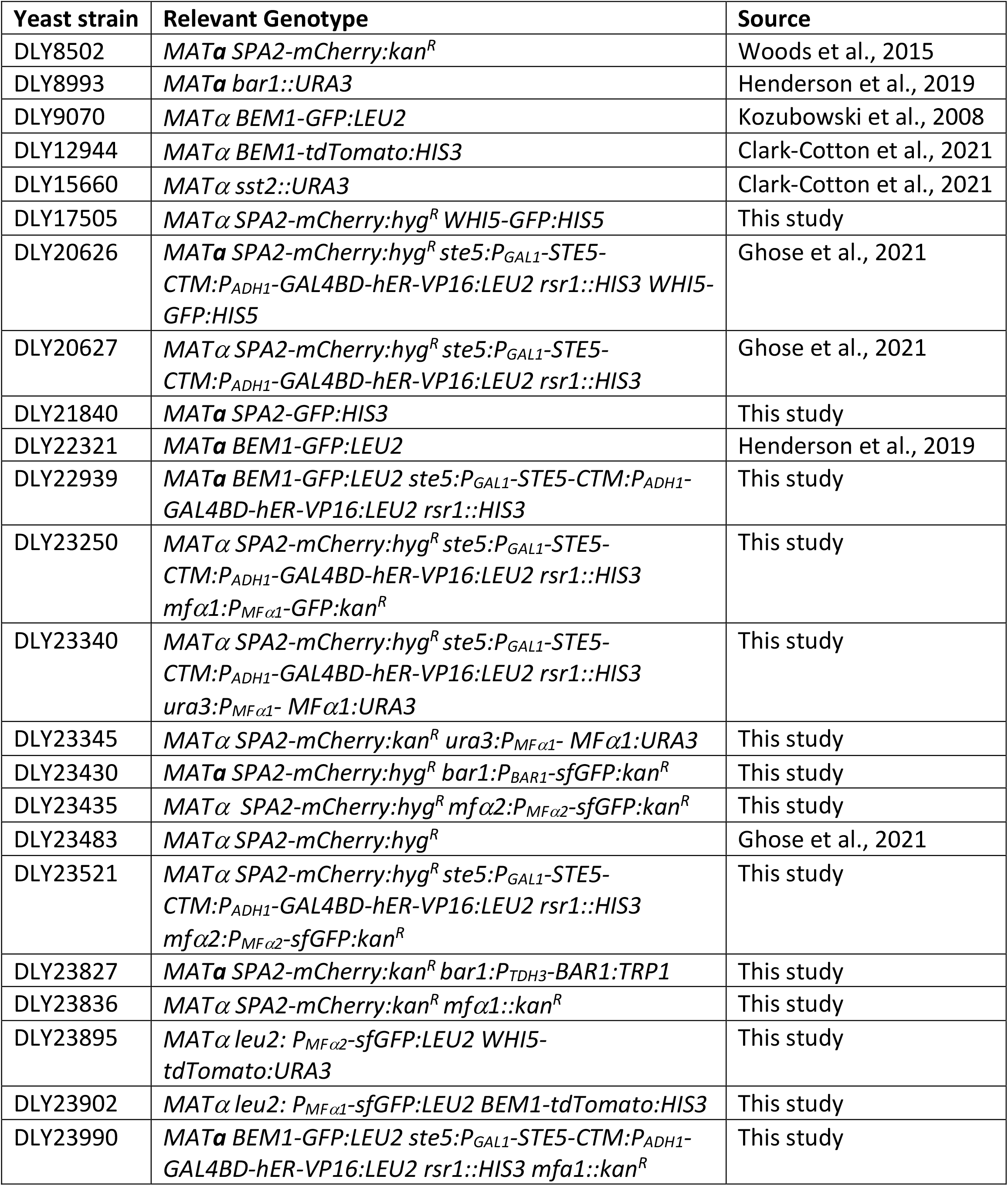
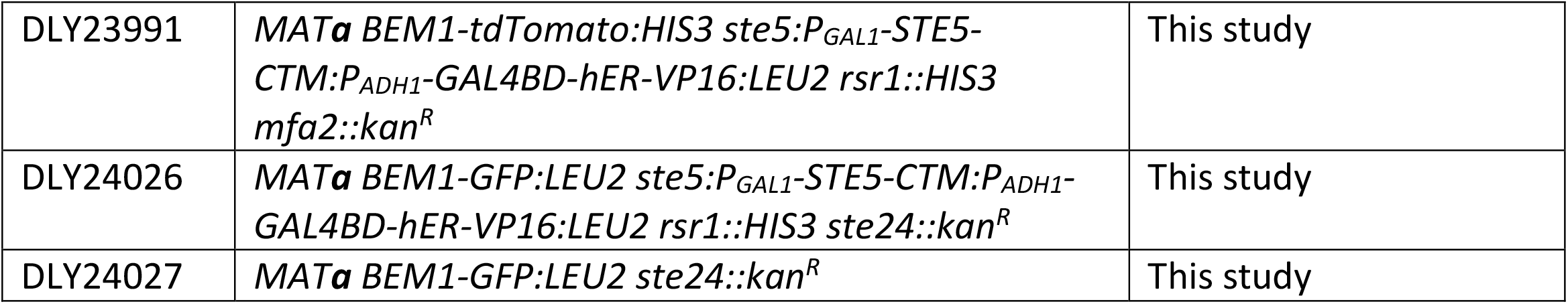

To delete *MFα1, MFα2, MFa1, MFa2, BAR1* and *STE24* we used a PCR-based method (Baudin et al., 1993) to precisely replace the open reading frame of the gene with heterologous sequences. Briefly, we used primers with 50 bp of homology to the 5’ and 3’ untranslated regions of the gene to be deleted to amplify a selectable marker from a template, transformed the PCR product into yeast, selected for the marker, and checked correct integration by PCR using flanking DNA primers. The templates were pFA6a-kanMX6 (*MFα1, MFa1, MFa2, STE24*) (Longtine et al., 1998) and pFA6a-link-yoSuperfolderGFP-kanMX6 (*MFα2, BAR1*) (gift from Wendell Lim & Kurt Thorn, Addgene #44901, (Lee et al., 2013)).

In DLY23250, we used the PCR-based method to replace the *MFα1* ORF with GFP by using genomic DNA from DLY20922 (McClure et al., 2018) as template.

To generate strains with an extra copy of *MFα1*, we PCR-amplified the *MFα1* ORF flanked by 500 bp upstream and 382 bp downstream from YEF473 genomic DNA with primers that added *ApaI* and *BamHI* sites, and cloned the fragment into the corresponding sites in a *URA3*-marked integrating plasmid (pRSII306) to make DLB4469. DLB4469 was digested with *NcoI* to target integration at *ura3*.

To replace the endogenous *BAR1* promoter with the *TDH3* promoter, the *TDH3* promoter was PCR-amplified from p405-TDH3-YFP (gift from Nicolas Buchler) with primers that added *BglII* and *PacI* sites and cloned the promoter into the corresponding sites in pFA6a-TRP1-PGAL1-GFP (Longtine et al., 1998)(replacing the GAL1 promoter) to make DLB4556. We then used the PCR-based method with DLB4556 as template to replace the endogenous *BAR1* promoter with the *TDH3* promoter.

To a generate a transcriptional reporter for *MFα2*, we PCR-amplified the promoter (500 bp upstream of the ATG) from YEF473 genomic DNA with primers that added *ApaI* and *HindIII* sites and cloned it into the corresponding sites in pDLB4448, upstream of the sfGFP. We digested the resulting plasmid (DLB4580) with *ApaI* and *SacI* to obtain P_MF*α*2_-sfGFP and cloned this fragment into a *LEU2*-marked single integration vector, SIVl (gift from Serge Pelet, Addgene #81090, (Wosika et al., 2016)), to make DLB4582. DLB4582 was digested with *BstBI* to target integration at *leu2*.

To generate a transcriptional reporter for *MFα1*, we PCR-amplified the promoter (500 bp upstream of the ATG) from YEF473 genomic DNA with primers that added *ApaI* and *HindIII* sites and cloned it into the corresponding sites in DLB4582 to replace P_MFα2_ with P_MFα1,_ generating DLB4600. DLB4600 was digested with *BstBI* to target integration at *leu2*.

### Live cell microscopy

Cells were grown overnight at 30°C in complete synthetic medium (CSM, MP Biomedicals) with 2% dextrose to mid-log phase (10^6^-10^7^ cells/mL). For mating mixtures, strains were mixed to obtain a 1:1 ratio unless otherwise indicated. Cell mixes were mounted on CSM slabs with 2% dextrose solidified with 2% agarose and sealed with petroleum jelly. For Ste5-CTM induction, cells were pre-treated with 20 nM β-estradiol (Sigma-Aldrich) for 3 h and mounted on slabs containing 20 nM β-estradiol. Imaging was performed at 30°C.

Images were acquired with an Andor Revolution XD spinning disk confocal microscope (Andor Technology) with a CSU-X1 5,000-rpm confocal scanner unit (Yokogawa) and a UPLSAPO 100x/1.4 oil-immersion objective (Olympus), controlled by MetaMorph software (Molecular Devices). Images were captured by an iXon 897 EMCCD camera (Andor Technology) or an iXon Life 888 EMCCD camera (Andor Technology).

Z-stacks with 15 z-steps of 0.48 *μ*m were acquired at 2 or 4 min intervals. Laser power was set to 10% of maximum for 488 nm and 13% of maximum for 561 nm. Exposure time was 250 ms and EM gain was 200.

### Halo assays to measure pheromone secretion

Pheromone from one mating type causes cells of the opposite mating type to arrest in G1. In the Halo Assay, a lawn (10^5^ cells) of supersensitive cells is spread evenly on a YEPD (1% yeast extract, 2% peptone, 2% dextrose) plate. A spot of cells (2 ⨯ 10^6^ cells) of the opposite mating type is pipetted onto the lawn, and pheromone diffusing away from the spot promotes G1 arrest, creating a zone of growth inhibition (the halo) in the lawn whose area reflects the amount of pheromone secreted from the spot. Supersensitive strains were DLY8993 (MAT**a** *bar1Δ* to prevent α-factor degradation) to detect α-factor and DLY15660 (MATα *sst2*Δ) to detect **a**-factor.

Because pheromone sensing and consequent MAPK activation leads to an increase in pheromone secretion, the amount of pheromone made depends on whether the secreting strain has active MAPK. To assess MAPK-induced levels of pheromone secretion, we spotted cells that contained inducible Ste5-CTM, and induced MAPK activation by including 100 nM β-estradiol on the assay plates. Strains to be tested and strains used as lawns were grown overnight at 30°C in YEPD to mid-log phase before spreading (lawn) or spotting. For **a**-factor halos, plates contained 0.04% Triton X-100, which enhances the spread of prenylated pheromone (Huyer et al., 2006). Halos were imaged after 48 h at 30°C. Halo area was measured by fitting a circle to each halo in ImageJ.

### Scoring commitment

In our mating mixes, there can be trivial reasons why a cell fails to commit during imaging. For example, a cell may not touch any cells of the opposite mating type (isolation), or its potential partners may be in the wrong stage of the cell cycle, or a potential partner may commit to a different cell. In these cases the failure to commit is uninformative. Therefore, only cells that had an opportunity to commit were scored: cells touching an opposite mating type partner that entered G1 early enough to allow time for commitment during the experiment (within the first hour of a two-hour time-lapse). Cells whose partners committed to another cell (a competitor) were excluded. Eligible cells were scored into three categories: (i) committed to a mating partner (defined by having a polarity site stably oriented toward a mating partner for at least 10 min), (ii) budded (failure of G1 arrest), or (iii) arrested in G1 but without achieving commitment during the imaging interval (these cells might eventually have gone on to commit or to bud). The percentage of cells in each category was calculated.

### Measuring indecisive and committed phases

The indecisive phase was defined as the interval between birth (cytokinesis) and commitment. The committed phase was defined as the interval between commitment and fusion. Cytokinesis was recorded at the time point when there was strong Bem1 or Spa2 signal at the bud neck. Commitment was recorded at the time point when the polarity site reached its final stable location and the Bem1 or Spa2 signal increased in intensity. Fusion was recorded at the time point when cytoplasmic mixing occurred (detected by the fluorescent probes in each mating type).

### Measuring *MFα1* and *MFα2* reporter expression

GFP fluorescence of the *MFα1* and *MFα2* reporters was measured at the time of commitment (scored based on Spa2 signal in the MAT**a** mating partner). On a maximum projection of the z-stack, an ROI was drawn manually around each cell in ImageJ and the mean fluorescence value was recorded. The mean fluorescence for an ROI where no cells were present was recorded as background, and the background value was subtracted from the cell fluorescence. The cell fluorescence values were normalized such that the mean of the *MFα1* reporter was set to 0.8 and the mean of *MFα2* reporter was set to 0.2 (to reflect the fact that *MFα1* and *MFα2* produce 80% and 20% of mature α-factor, respectively (Rogers et al., 2012)).

### Measuring cell area

Cell area was measured at the time of cell birth (cytokinesis scored by Spa2 signal at the bud neck). An ROI was drawn manually around each cell in ImageJ and the area was recorded.

### Determining MATα:MATa ratio

When mating types were mixed at a ratio other than 1:1, the final ratio was empirically determined by counting the number of MATα and MAT**a** cells in each of 8 stage positions. The ratio of MATα:MAT**a** cells in each stage position was calculated and the values were averaged to get a MATα:MAT**a** ratio for the experiment; error bars represent standard deviation.

### Scoring polarity site encounters

Polarity sites in cells expressing Ste5-CTM were analyzed using maximum projections of z-stacks in ImageJ. An encounter was recorded when the polarity sites (visualized using Spa2 probes) in partner cells approached within 1 µm of each other. Images were visually inspected to identify times with potential encounters. A line 10 pixels wide was drawn across the two polarity sites to generate a line scan of fluorescence intensity. Each polarity site appeared as a peak of signal intensity on the line scan and the distance between the two peaks was measured. An encounter was recorded if peaks were less than 1 µm apart.

### Statistical analysis

Two-sample Kolmogorov-Smirnov (KS) tests and T tests were performed in Microsoft Excel using the Real Statistics Resource Pack add-in (Release 5.11) developed by Charles Zaiontz.

## Data availability

Experimental data used to generate figures are included in the S1 Data spreadsheet.

## Acknowledgements

We thank Beverly Errede, Nick Buchler, Tim Elston and members of the Lew lab for helpful conversations and comments on the manuscript. We thank Yasheng Gao and Lisa Cameron of the Duke Light Microscopy Core Facility for assistance with microscopy. This work was funded by National Institutes of Health/National Institute of General Medical Sciences Grant no. R35GM-122488 to D.J.L.

## Notes

### Competing Interest Statement

The authors have declared no competing interest.

